# Nanostructured lipid carriers overcome the low immunogenicity of M2e peptide via surface click chemistry conjugation, improving the anti-M2e antibody response

**DOI:** 10.1101/2025.08.25.672124

**Authors:** Louis Bourlon, Carole Fournier, Antoine Hoang, Christine Charrat, Alban Lepetitcolin, Jessica Morlieras, Fabrice Navarro, Patrice N Marche

**Affiliations:** Univ. Grenoble Alpes, Inserm U 1209, CNRS UMR 5309, Institute for Advanced Biosciences, Grenoble, France; Microtechnologies for Living Systems Interactions Research Unit, Univ. Grenoble Alpes, CEA, LETI, Technologies for Healthcare and Biology Division, Grenoble, France; Sanofi, Research and Development, Marcy L’Etoile, France

**Keywords:** Nanoparticles, Nanostructured Lipid Carriers (NLC), Matrix 2 Ectodomain (M2e), Click chemistry, Vaccine, Influenza

## Abstract

Influenza vaccines are considered the most effective measure for preventing influenza. The key antigen of commercial vaccines is hemagglutinin (HA), the main protein on the surface of influenza viruses. This protein is genetically variable, necessitating the annual redefinition of the vaccine formulation. Furthermore, these vaccines do not offer protection against pandemic viral strains. The M2e peptide, a small surface protein conserved among influenza type A strains, could enable cross-immunity when used as an antigen. Nevertheless, its limited immunogenicity has restricted its application in universal influenza vaccines. To enhance the immune responses to M2e, we associated it with Nanostructured Lipid Carriers (NLCs). These NLCs are stable over time and their small diameter (<100 nm) enhances interactions with antigen-presenting cells and improve lymphatic drainage, facilitating their uptake by the immune system. The M2e specific antibody response in mice was enhanced by using “click” chemistry to conjugate the M2e peptide to the NLC surface. Also, the incorporation of PADRE-specific CD4+ T cells stimulating peptide between M2e peptide and its linking region greatly enhanced the specific antibody response to M2e and generated immune response levels that could be compatible with influenza protection. This promising M2e formulation vectorized by NLC represents a potential universal influenza vaccine candidate warranting further preclinical investigations and clinical evaluation.

## INTRO

Seasonal influenza remains a significant global health burden, causing an estimated 3-5 million severe cases and up to 650,000 respiratory-related deaths annually [1]. The influenza virus is notorious for its ability to mutate rapidly, leading to antigenic drift and shift, which necessitates annual reformulation of vaccines to match circulating strains. While current vaccines are effective in protecting against seasonal strains, they fail against emerging pandemic strains, as demonstrated during past outbreaks such as the 2009 H1N1 pandemic. The unpredictable nature of the influenza virus evolution underscores the urgent need for a universal influenza vaccine capable of providing broad, lasting protection against both seasonal and pandemic strains [2].

In this context, the M2e peptide, the conserved extracellular domain of the influenza virus matrix protein 2, has emerged as a promising candidate for a universal vaccine. Unlike more variable antigens like hemagglutinin (HA) and neuraminidase (NA), M2e remains relatively conserved across influenza A subtypes. However, M2e is poorly immunogenic by itself possibly due to its small size and low abundance on the viral surface compared to HA and NA [3]. This limited immunogenicity raises a challenge for eliciting a robust immune response against M2e when it is included in conventional vaccine formulations. To address this limitation, vaccine platforms must be designed to effectively deliver M2e and enhance its immunogenic properties. Various M2e-based constructs have been developed using different delivery strategies and have shown protective efficacy *in vivo* in both mice and ferrets [4]. However, the only Phase I clinical trial conducted in humans using an M2e-VLP vaccine produced disappointing results, with a rapid decline in anti-M2e IgG titers over time. These findings underscore the need for innovative vectorization platforms to enhance the immunogenicity and durability of M2e-based vaccines [5].

Lipid nanoparticles (LNPs) represent a versatile platform for vaccine delivery, enabling encapsulation or conjugation of antigens, adjuvants, and other immunomodulatory molecules. [6]. Among these, a new generation of LNPs named nanostructured lipid carriers (NLCs) possess a solid core and offer several advantages, including excellent stability and biocompatibility [7]. NLCs can be tailored to modulate antigen presentation and can present antigens in a multimeric format, which enhances B cell recognition and antibody production [8]. Moreover, NLCs can be engineered to co-deliver adjuvants, such as CpG oligodeoxynucleotides. These adjuvants play a pivotal role in promoting a strong T helper 1 (Th1)-biased immune response essential for cytotoxic T cell activation and the overall effectiveness of vaccine-induced immunity [9] [10]. The Th1 response is characterized by the production of pro-inflammatory cytokines such as interferon-gamma (IFNγ), which not only enhances the activation of various immune cells but also supports antibody class switching of antibodies towards more effective isotypes like IgG2a, improving opsonization and clearance of infected cells [11] [12].

In this study, we investigated the use of NLCs as a delivery platform for M2e peptide, with the goal of enhancing the immunogenicity of this otherwise weak antigen. Click chemistry (strain-promoted azide-alkyne cycloaddition (SPAAC)) [13] offered a powerful method to conjugate M2e to NLCs ensuring the preservation of crucial epitopes and optimizing antigen density and presentation for improved immune recognition. Additionally, we explored the inclusion of a flexible linker composed of five amino acids between the M2e and the NLC, as well as the inclusion of the universal helper T cell epitope PADRE (PAN-DR epitope) [14] to further amplify the anti-M2e immune response.

The immune response triggered by our NLC-M2e formulation generates immune response levels that are compatible with influenza protection, even if further clinical evaluations would be required to formally access the vaccination. Globally, our findings provide important insights into how NLC-based vaccines can be optimized for use in universal influenza vaccines and other immunologically challenging targets.

## EXPERIMENTAL SECTION

### Materials

Suppocire NC™ was purchased from Gattefossé (France). Myrj™ S40, polyethylene glycol (PEG) 40 stearate, and Super Refined Soybean Oil came from Croda Uniqema (Chocques, France). SA-PEG100-NH2 was provided by Eras-Labo (France) and Dibenzocyclooctyne-N-hydroxysuccinimidyl ester (DBCO-NHS ester) by Click chemistry tool (China). Lipoid S75 (soybean lecithin at >75% phosphatidylcholine) was purchased from Lipoid (Germany). 1,2-dioleoyl-3-trimethylammonium-propane chloride (DOTAP) and 1,2-Dioleoyl-sn-glycero-3-phosphoethanolamine (DOPE) was provided by Avanti Polar Lipids (USA). 1,1’-Dioctadecyl-3,3,3’,3’-Tetramethylindodicarbocyanine perchlorate DiD (emitting at 565 nm) was purchased from Invitrogen (France). 2-azidoethanol was provided by (Thermo Fisher) Other chemical products came from Sigma-Aldrich (France), including N,N-Diisopropylethylamine (DIEA) and Dichloromethane (DCM). The free M2e peptide was ordered from MedChemExpress and stored at 10 mg mL^-1^ in pure DMSO at −20°C. Various modified M2e peptides were synthesized by Provepharm (detailed sequence in Table S1). Peptides were stored lyophilized at −20°C for long-term preservation and resuspended at 10 mg mL^-1^ in pure DMSO for solubilization and immediate use.

### DBCO-modified-lipid synthesis: SA-PEG100-DBCO

To synthesize SA-PEG100-DBCO, SA-PEG100-NH2 (38 µmol) of DIEA (0,3mmol) were dissolved in DCM (10mL) under inert atmosphere. After 5 minutes of stirring, DBCO-NHS ester (76 µmol) (Click chemistry tool) were added and the whole solution was stirred for 1h. The reaction was monitored by thin layer chromatography (TLC) (CH2Cl2/MeOH 9/1). The final product was precipitated twice in ether and then dialyzed against milli-Q water (cut-off threshold 1000 Da, ZelluTrans). Water was removed by freeze-drying. Purity was checked by 1H NMR analysis (∼100% of SA-PEG100-NH2 is converted in SA-PEG100-DBCO).

### Formulation of neutral NLCs

The lipid phase and the aqueous phase were prepared separately. The lipid phase was prepared by dissolving wax, soybean oil and lecithin S75 in ethanol at RT. When needed, fluorescent NLC were used for quantitative characterizations. For this purpose, a lipophilic dye solution (DiD) in EtOH was added to the lipid phase. Ethanol was evaporated at 47°C under nitrogen for approximately 15 minutes. Ethanol was evaporated at 47°C under nitrogen for approximately 15 minutes. The aqueous phase was prepared by dissolving both unmodified and modified PEGylated lipids, respectively SA-PEG40-OH and SA-PEG100-DBCO in PBS1x. Different modified/unmodified PEGylated lipids ratios were tested, thus leading to different DBCO moieties densities: final formulations displayed either 1.6% or 5% (w:w) modifiable DBCO moieties at the NLC surface. The aqueous phase was added to the lipid phase and the final solution was mixed through high frequency sonication. NLCs were purified by dialysis against 1,000 volumes of PBS1x and stored at 4°C [9].

### Formulation of cationic NLCs

The lipid phase and the aqueous phase were prepared separately. The lipid phase was prepared by dissolving wax, soybean oil, lecithin SCP3, DOTAP and DOPE in ethanol at RT. Ethanol was evaporated at 47°C under nitrogen for approximately 15 minutes. Ethanol was evaporated at 47°C under nitrogen for approximately 15 minutes. The aqueous phase was prepared by dissolving the unmodified PEGylated lipid (SA-PEG40-OH) in PBS1x. The aqueous phase was added to the lipid phase and the final solution was mixed through high frequency sonication. NLCs were purified by dialysis against 1,000 volumes of PBS1x and stored at 4°C [15].

### Physicochemical characterization

Dynamic Light Scattering (DLS)measurements were performed using the ZetaSizer Nano ZS (Malvern Instruments) and used to determine the hydrodynamic diameter of NLCs. Measurements were conducted in a 1x PBS solution, pH 7.4, at room temperature, with a NLC concentration of 1 mg mL^-1^. Each analysis consisted of three independent measurements. Each measurement comprised 10 repetitions of 10 seconds each. Measurements were performed at a fixed angle of 173° using a 633 nm laser. Data analysis was performed using the cumulant analysis algorithm.

Nanoparticle tracking analysis (NTA) measurements were performed using the NS300 NanoSight (Malvern Instruments) with a 488 nm laser. The sample was systematically diluted by a factor of 5 × 10^5^ or 1.5 × 10^5^, depending on the expected concentration, to ensure it remained within the optimal reading range of the NS300. Each measurement consisted of three repetitions, with a recording time of 1 minute per repetition. To account for measurement and dilution variability, three separate measurements were conducted with three different dilutions. The average of these three measurements was used for analysis. The injection was performed via a syringe pump at 25°C.

### Peptide conjugation to NLCs

The peptide of interest was first dissolved in pure dimethyl sulfoxide (DMSO) at 10 mg mL^-1^ and was then carefully diluted in 10 volumes of water to reach 1 mg mL^-1^. The peptide was then added dropwise to the NLC-DBCO solution (PBS1x at pH 7.3) at room temperature at different molar ratios depending on the desired final peptide density (Table S2). The reaction was gently stirred for different durations (Table S2 for optimized durations). Remaining DBCO groups were capped by 2-azidoethanol (molar ratio 20 eq. N_3_:1 eq. DBCO) for 15 minutes at room temperature.

### NLC-M2e Purification by size exclusion chromatography

At the end of the reaction, the solution was purified by size exclusion chromatography (SEC) using Superdex 200 resin (Cytiva – 28990944 ; Fractionation range from 10 000 to 600 000 M_r_ (Molecular mass), Exclusion range at 1 300 000 M_r_). The process was controlled by an AKTA Pure 25 system (Cytiva). Elution was performed at a constant flow rate of 0.75 mL min^-1^ with 1x PBS. The mixture was introduced into the system by aspiration with automatic air detection. The solution was diluted to an NLC concentration of < 4% (w/v) to avoid saturation of the column. Absorbance at 280 nm and conductivity were monitored to estimate the fractions containing the peptide and/or the NLC. Fractions were collected using the F9C fraction collector (Cytiva). The proportion of collected NLCs was estimated by summing the absorbance at 280 nm of the collected fractions relative to the total absorbance of all NLC-containing fractions. The collected fractions were then combined into a single solution.

### Conjugation rate estimation

The conjugation efficiency of the peptide to NLC-DBCO corresponds to the percentage of peptide successfully conjugated to NLC-DBCO relative to the initial amount of peptide introduced. This efficiency can be estimated using fluorescently labeled versions of the peptide and the NLC: A488-M2e-N_3_, *i.e*., the peptide labeled at the N-terminus with the fluorophore Alexa Fluor 488 (Excitation/Emission: 490/525 nm); NLC(DiD)-DBCO, *i.e.*, an NLC functionalized with DBCO and incorporating the lipophilic fluorophore DiD in its core (Excitation/Emission: 644/655 nm).

The A488-M2e-N_3_ peptide is added to the NLC(DiD)-DBCO in aqueous solution, and the resulting mixture is purified by size exclusion chromatography as described above. Fluorescence is then measured in both the A488 and DiD channels for each collected fraction. NLC(DiD)-M2e conjugates, having a larger hydrodynamic volume than free A488-peptides, elute in the early fractions and are detected as a single peak based on DiD fluorescence. Free A488-peptides elute later and are quantified via A488 fluorescence. In cases of effective conjugation, two A488 fluorescence peaks are observed: one coinciding with the DiD-labeled NLC peak (corresponding to conjugated peptides), and a second, distinct peak (corresponding to unbound peptides).

The conjugation efficiency is calculated by integrating the area under the respective A488 fluorescence peaks, using the following formula:

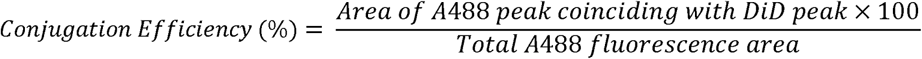

### Peptide concentration of the NLC-peptide solution

The peptide concentration in the conjugated solution was determined by Amino Acid Analysis (AAA) using Ultra Performance Liquid Chromatography (UPLC). This approach is more direct and less prone to bias than the conjugation rate method (described above), and is therefore preferred for accurate quantification of peptide concentration. The solution containing NLC mixed with peptides was diluted in PBS 1x to achieve an NLC concentration below 1% (w/v) and a volume of 100 µL. Each sample was run in duplicate. Samples were hydrolyzed in 6N HCl heated to 110°C for 16 hours in closed tubes. At the end of the reaction, the temperature was lowered to 80°C, the tubes were opened, and HCl was evaporated under an argon stream. The hydrolysate was resuspended in 20 mM HCl (200µL) and derivatized with the fluorescent compound AccQTAG (Waters). UPLC was performed using a reverse phase column (Waters).

### M2e density calculation

The approximate density of M2e per NLC can be expressed in different and complementary ways.

- In quantity of M2e per NLC, calculated as the molar ratio N(M2e)/N(NLC), with N(M2e) being obtained by AAA-UPLC and N(NLC) being obtained by NTA.
- In quantity of M2e per surface area: 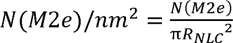
- If M2e is considered to be evenly distributed on the NLC, with the sphere being equivalent to a flattened surface, the distance between each M2e can be approximated by this distance:

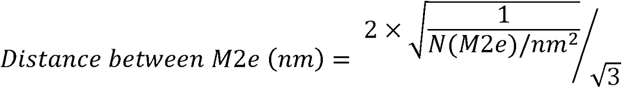

### Animals

Wild type female BALB/c mice were supplied by Charles River and housed in the animal facility at the Institute for Advanced Biosciences (La Tronche, France) with a maximum of 6 mice per cage. The number of mice in each treatment group, for each experiment, is indicated in the legend of the figures. The primary outcome measure was the M2e specific antibody measurement and this parameter was used for the sample size calculation (6 for experimental conditions, 4 for negative controls). Animal use and care were approved by the Animal Experiments Ethics Committee “Comité d’Ethique pour l’Expérimentation Animale no.#12, Cometh-Grenoble” and approved by the French Ministry of Research (#26785-2020073017416031 v2). Based on this document, no mice were excluded based on preestablished criteria. Immunization and analysis protocols were designed to avoid expected cofounders. In particular, all mice comparisons in each figure are from the same experiment (ie, same timing of immunization and analysis), except otherwise stated. L.B was aware of the group allocation during all the stages of the experiments. All experiments were performed in accordance with ARRIVE guidelines.

### Immunizations

Mice were immunized intraperitoneally with an initial injection at the age of 8-9 weeks followed with two booster shots on days 21 and 28. Peptide and NLC-peptide doses are indicated in each concerned figure (from 0.6 to 4.4µg per injection). CpG ODN 1826 (10 µg per injection; InvivoGen, #tlrl-1826) was used as an adjuvant. The CpG was eventually vectorized by NLC(+) (NLC(+)/CpG), as documented in the corresponding figure. One week after the last immunization, blood samples were collected, the mice were humanly euthanized, and spleens were collected.

### Splenocyte restimulation

Splenocytes were extracted by mechanical grinding through a 100 µm cell strainer and collected in RPMI medium (Gibco 61870–010). The cell suspension was centrifuged for 5 min at 350 g, and erythrocytes were lysed using 1 mL of RBC lysis buffer (Biolegend 420302). Cells were washed by centrifugation in 10 mL RPMI medium and resuspended in RPMI medium supplemented with 10% FBS, 1% non-essential amino acids (PAA M11-003), 1% penicillin-streptomycin (Invitrogen), and 1% sodium pyruvate (GE Healthcare S11-003). Cells were then seeded at 10^6^ cells/mL in 24-well plates (500 µL/well) and stimulated with 2 µg of free M2e. For each sample, a negative control with unstimulated cells and a positive control with CD3/CD28 activation beads (Gibco, 11456D) were prepared. After 2 days of incubation at 37°C, plates were centrifuged for 5 min at 350 g to collect the supernatant which was stored at −20°C. To analyze IFNγ and TNFα produced by splenocytes, a protein transport inhibitor (eBioscience™ Protein Transport Inhibitor Cocktail, Invitrogen, 00-4980-93) was added to the culture medium 16 hours before analysis by flow cytometry.

### Antibody Assay by ELISA

ELISA was performed in a MaxiSorp® 96-well plate (Thermofisher). All steps were carried out under agitation, except for the M2e incubation. Each step was followed by three washes with 300 µL of PBS 1x Tween 0.05%, using an automatic plate washer (WellwashTM, Thermofisher). 100 µL of M2e (1 µg mL^-1^, diluted in PBS 1x) were added per well and incubated at 4°C in the dark for 16 hours. The isoelectric point (pI ∼4) of M2e allowed it to adhere to the plate at this pH (7.4). The plate was neutralized by adding 200 µL of blocking buffer (ELISA/ELISPOT diluent 1x, Thermofisher, 00-4202) and incubated for 1 hour at room temperature. The standard curve of the influenza A m2 antibody 14C2 (Santa Cruz Biotechnology, sc-32238) was prepared by two-fold serial dilution, starting at 10 µg mL^-1^ down to 9.8 ng mL^-1^. Meanwhile, sera were diluted in blocking buffer according to the expected antibody quantity, with a minimum dilution of 1:10, and incubated for 2 hours at room temperature. Detection was performed by adding an anti-IgG-peroxidase antibody (SigmaAldrich - A9044) diluted 1:40,000 in blocking buffer and incubated for 1 hour at room temperature. Wells were then washed six times to ensure no unbound anti-IgG-peroxidase antibody remained. 100 µL of TMB solution (3,3’,5,5’-Tetramethylbenzidine, Invitrogen, 002023) were added and incubated at room temperature in the dark for 10 minutes. The reaction was stopped by adding 50 µL of 2N H2SO4.

To perform ELISA(NLC-M2e), 100 µL of the NLC solution at 40 µg mL^-1^ were placed in a MaxiSorp® 96-well plate. The plate was incubated at 4°C for 16 hours in the dark. The protocol then followed the same steps as above. The NLCs were revealed using 14C2 at 1 mg mL^-1^.

### ELISA (Cell-M2)

A549 cells human lung epithelial adherent cells, were cultured in DMEM with 4.5 g L^-1^ glucose + Glutamax supplemented with 10% fetal bovine serum. M2 mRNA was custom-ordered from RiboPro with the specified coding sequence. The optimization of the 3’ and 5’ UTRs was performed by the supplier. mRNA was supplied in RNase-free water at a concentration of 1 mg mL^-1^, aliquoted, and stored at −80°C for single use.

For transfection, A549 cells between passages 3 and 7 were spread in Opti-MEM medium (Gibco 31985) and cultured in a 96-well plate. Each well contained 5×10^4^ cells, optimized to obtain a sufficient signal during ELISA. The cells were cultured at 37°C; 5% CO_2_ overnight. The next day, cells were transfected with M2 mRNA using Lipofectamine™ MessengerMAX™ transfection reagent (Invitrogen, LMRNA001) according to the supplier’s instructions. Six hours after transfection, the medium was changed to DMEM with 4.5 g L^-1^ glucose + Glutamax + 10% fetal bovine serum. Transfection continued for an additional 18 hours at 37°C; 5% CO2.

The next day, 100 µL of 8% formaldehyde was added to each well for 15 minutes at room temperature. The plate was then washed 6 times with PBS 1x. The rest of the protocol was identical to the one described earlier, in the “Antibody Assay by ELISA (M2e)” section. Fixed cells could be stored for at least one week at 4°C in the dark without affecting detection. Non-specific antibody binding was observed, although significantly lower than specific binding to M2e. This non-specific binding was proportional to the immune response. Each serum was tested in parallel on a non-transfected cell, with the non-specific OD from non-transfected A549 subtracted from the specific OD obtained with A549-M2.

### Competitive Inhibition ELISA (Cell-M2)

Serum from immunized mice was mixed at a fixed dilution with increasing concentrations of free M2e peptide. The mixture was incubated for 1 hour at room temperature with agitation. The mixture was then assayed on the Cell-M2 with the well background saturated by “ELISA/ELISPOT 1x diluent.” The protocol followed the same steps as previously described.

### IFN**γ** Quantification by ELISA

This assay quantified IFNγ in the supernatant of restimulated splenocytes. It was conducted as per the supplier’s instructions (“IFN gamma Mouse Uncoated ELISA Kit”; 88-7314-88; sensitivity: 15 pg mL^-1^).

### ELISA Analysis

Optical density (OD) was read at 450 nm and 570 nm with Clario Star Plus (BMG LABTECH). OD values >2 were excluded, and the lowest possible OD above the limit of quantification was considered in the calculations. The OD at 570 nm was subtracted from the OD at 450 nm to remove non-specific absorbance, as per the Thermofisher kit instructions. The average of the two blanks (without antibody addition) was subtracted from all wells. The standard curve using 14C2 was performed in duplicate. The mean and standard deviation of the curve were calculated, and concentration was transformed by Y = Log(X). In GraphPad, the standard curve was plotted using an “Asymmetric five parameters” model with “weight= 1/Y^2” and the constraint “Bottom Must be greater than 0.” Other parameters were left at their default values. When multiple dilutions yielded calculable results, the lowest dilution that produce an OD value at least three times higher than the negative controls was selected.

### Flow cytometry

After supernatant removal, splenocytes were rinsed with PBS. Then, 50 µL of “LIVE Far Red®” viability marker (1:1000 in PBS 1x) was added to the cells. The plate was incubated for 15 minutes at room temperature in the dark, and then rinsed with flow cytometry staining buffer (FCSB) (eBioscience, 00-4222-26). Extracellular antibodies (Table S3) were diluted in FCSB and applied to the splenocytes (50 µL). Cells were incubated for 20 minutes at 4°C in the dark and rinsed with FCSB. The intracellular staining kit (eBioscience FOXP3/transcription Factor Staining Buffer Set, 00-5523-00) was used according to the manufacturer’s protocol for cell fixation and permeabilization. Intracellular antibodies (Table S3) were diluted in permeabilization buffer and added to the splenocytes (50 µL). Cells were incubated for 30 min at room temperature in the dark, then rinsed twice with FCSB. Splenocytes were resuspended in 200 µL of FCSB and analyzed by flow cytometry (Attune NxT, Thermofisher). Compensation beads were used for each marker, and splenocytes were stained with “all markers minus one” (Fluorescence Minus One, FMO) to establish the positive threshold for analysis. Data were analyzed using the Attune™ NxT Software.

### Statistical Analysis

All statistical analyses were conducted using GraphPad software. Data were examined for normal or log-normal distributions. If 2 groups were normally distributed, a t-test was performed with α=0.05. If more than 2 groups were normally distributed, an ordinary one-way ANOVA test was performed, followed by a post-hoc Holm-Sidak test with α=0.05 if significant (p<0.05). If groups were log-normally distributed, values were log-transformed before applying the same tests. If data were neither normally nor log-normally distributed, Mann-Whitney or Kruskal Wallis tests were performed.

## RESULTS

### NLC-M2e are efficiently produced and characterized

To enhance their immunogenicity, M2e peptides were conjugated to NLCs which have been proven to be efficient for protein delivery [9, 10]. For this purpose, neutral NLCs were functionalized with DBCO moieties on their surface (NLC-DBCO, Figure S1) yielding an average hydrodynamic size of 73.6 ± 2.0 nm (measured by DLS, Table S2). The conjugation of M2e-N_3_ (M2e peptide with an azide (N_3_) moiety at the C-terminal end) to NLC-DBCO *via* a SPAAC reaction [16] led to the formation of a triazole bond, resulting in the covalent conjugation of the peptide (NLC-peptide) (Figure 1). Remaining DBCO groups were capped by adding 2-azidoethanol. Unconjugated peptides and excessive 2-azidoethanol were subsequently removed by SEC. Regardless of peptide density, NLC-M2e globally showed a slight increase in their average hydrodynamic diameter compared to NLC-DBCO (82.4 ± 7.8 nm, measured by DLS (Figure 2A; Table S2)) and a monodispersed population (average PdI = 0.192 ± 0.017, measured by DLS (Figure 2A; Table S2)). Nanoparticle tracking analysis confirmed a homogeneous population, without aggregates up to the maximal size assessed of 1000 nm (Figure S2A).

**Figure 1.**
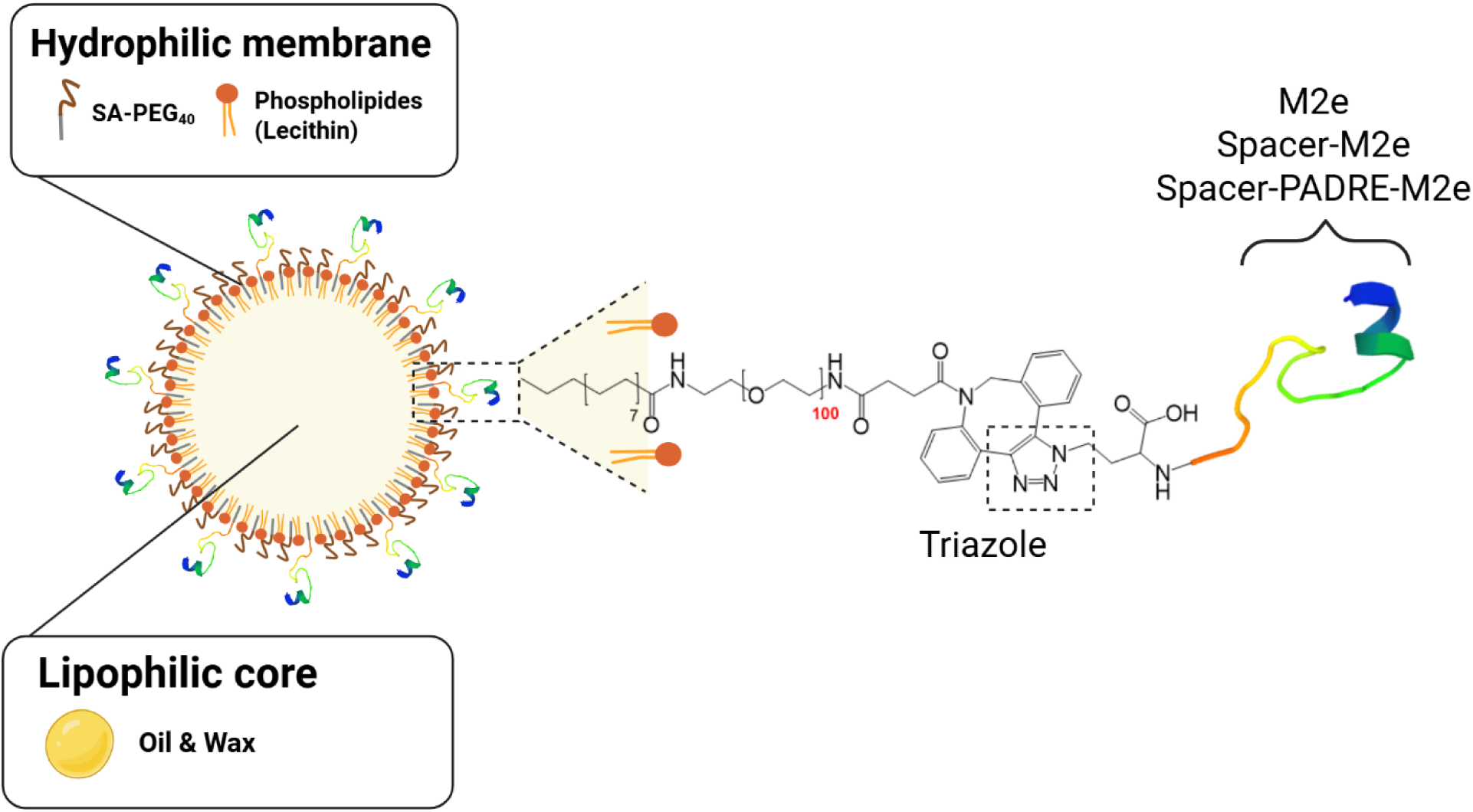
Schematic representation of NLC-peptide.

**Figure 2.**
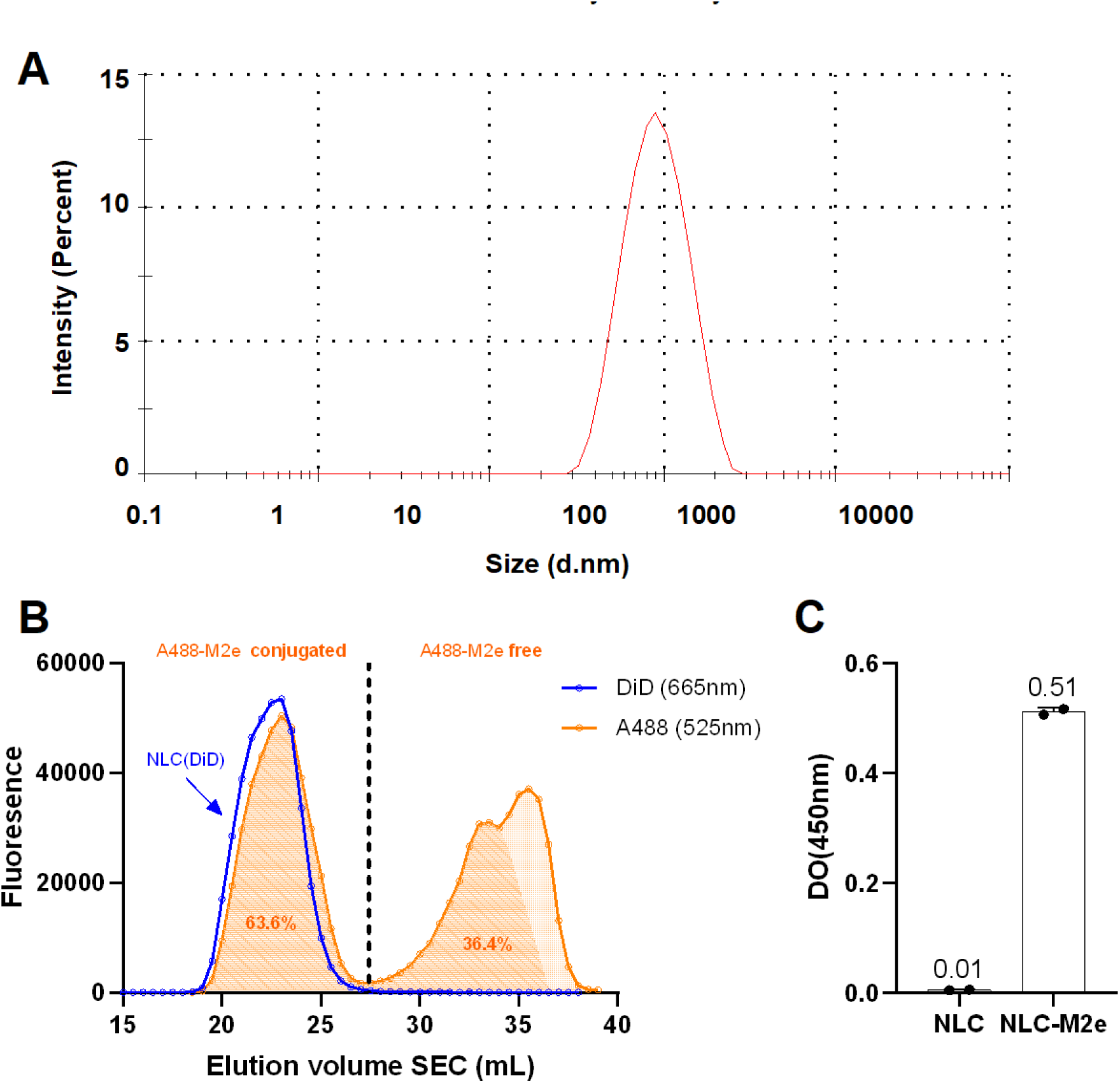
Conjugation of M2e with NLC. A Dynamic Light scattering (DLS) assessing the hydrodynamic size of the NLC-M2e at the day of the formulation. This graph is representative of a series of 7 measurements. yielding an average diameter of 82.7 ± 5.6 nm. B Estimation of the M2e conjugation to NLC-M2e by using fluorescent peptide and SEC. C M2e accessibility assessed by ELISA. using the anti-M2e commercial antibody 14C2.

Fluorescent M2e (A488-M2e-N_3_) and fluorescent NLC-DBCO (NLC(DiD)-DBCO) were used to estimate the conjugation efficiency as described in the Materials and Methods section.The NLC(DiD)-DBCO fraction was identified by DiD fluorescence as a single peak, eluted first (Figure 2B). Now, with regard to fluorescence at 488nm, the first peak, co-eluted with the NLC(DiD)-DBCO peak, represents the conjugated peptide, while the second peak corresponds to the free peptide. The unshaded region indicates free A488 fluorophore dissociated from the peptide (checked by AAA-UPLC, not shown). Integration of the shaded area enabled the calculation of the conjugation efficiency. Figure 2B illustrates an example of A488-M2e-N_3_ to NLC(DiD)-DBCO conjugation efficiency with an efficiency of 63.6%.

The conjugation efficiency also allowed for the indirect estimation of M2e molarity in the NLC-M2e solution, which was confirmed by AAA-UPLC analysis. Both methods demonstrated excellent correlation (Figure S2B) and minimal bias, as indicated by a Bland-Altman analysis (Figure S2C). The conjugation conditions were optimized to produce NLC-M2e with varying peptide densities, ranging from approximately 20 to 190 peptides per NLC (Table S2).

The accessibility of M2e on NLC-M2e was demonstrated by ELISA using anti-M2e antibody (14C2), which recognize M2e on the surface of NLC-M2e (OD= 0.52) but not on NLC alone (OD= 0.01), as shown in Figure 2C. These results confirm that M2e epitopes are accessible on the NLC-M2e surface.

Two additional peptides, namely “Spacer-M2e” to move M2e away from the NLC and potentially to help interaction with specific antibodies, and “Spacer-PADRE-M2e” to support specific immune response (Table S2), were also successfully conjugated (Table S2). All NLC-M2e and other NLC-peptide conjugates exhibited monodispersity with only minor variations in hydrodynamic size (Table S2), attributed to the nature of the peptide and/or the density.

Concerning the stability of the NLC-DBCO before conjugation to M2e-N_3_, the DBCO moieties maintained their reactivity over time, showing only a moderate reduction in conjugation efficiency (18.7%) for A488-M2e-N_3_ after 88 days in solution (Table S4).

### Focusing Anti-M2e Antibody Response through M2e Conjugation to NLC

To assess the efficacy of NLC-M2e in inducing anti-M2e immune response and to evaluate the advantages over free M2e, mice were immunized as described in M&M. The anti-M2e antibody response was assessed in the blood serum by two ELISA: one using M2e peptide as antigenic target (“ELISA(M2e)”) and the other using A549 cells expressing M2 protein (“ELISA(cM2)”).

The data presented in Figure 3 show that mice immunized with free M2e produced stronger anti-M2e IgG responses compared to mice immunized with NLC-M2e (average OD of 1.3 vs. 0.16, respectively, p=0.04, Figure 3A). Furthermore, immunization with either NLC-M2e or free M2e induced similar levels of anti-(NLC-M2e) antibodies (Figure S3). This indicates that NLC-M2e do not generate stronger antibody responses than free M2e.

**Figure 3.**
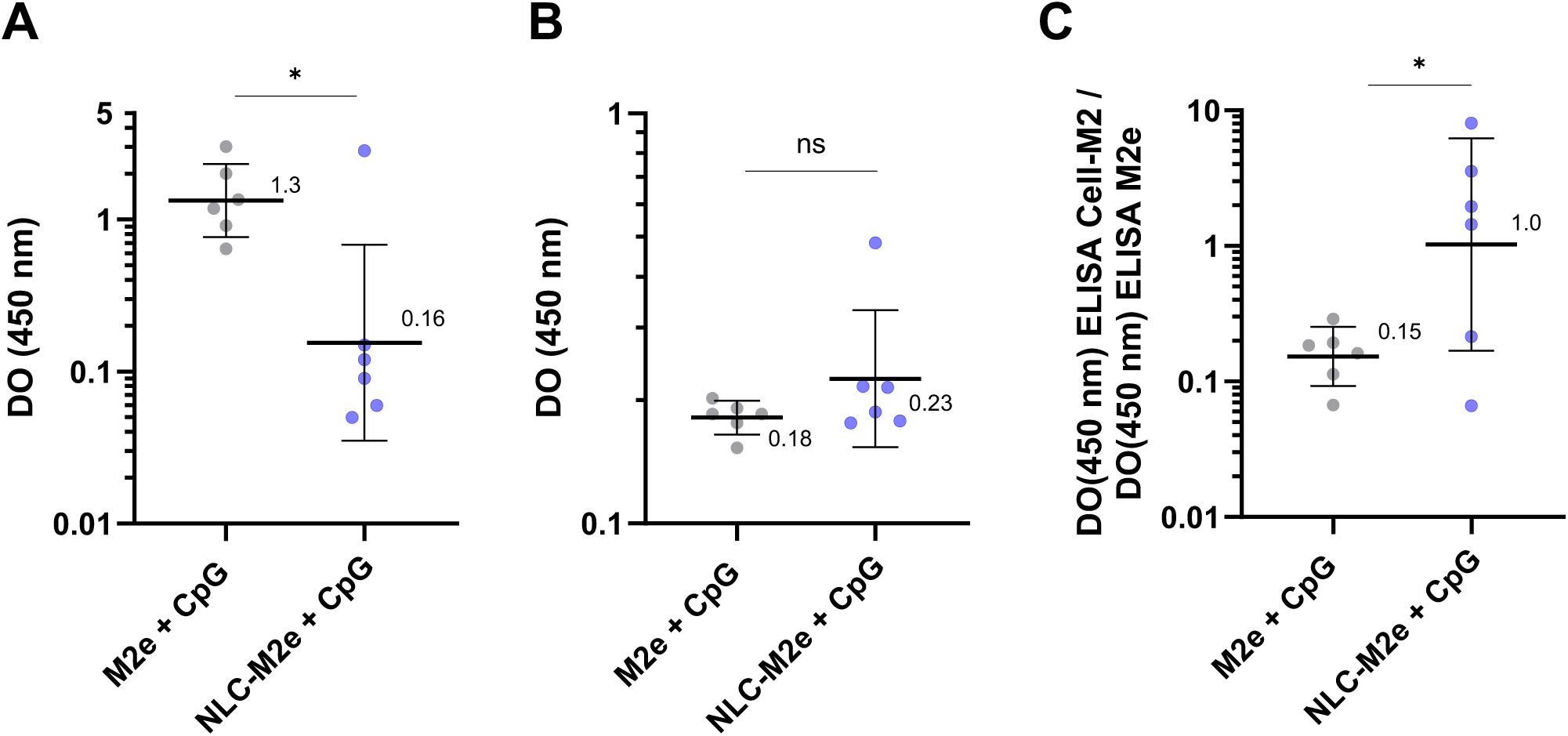
M2e vectorization through NLC-M2e allows to focus the antibody response against the native M2. BALB/c mice were immunized and double boosted with non-vectorized M2e (M2e + CpG) or vectorized M2e (M2e NLC-M2e + CpG) with 4µg M2e and 10µg CpG (ODN 1826). M2e specific antibodies in the blood serum were quantified by **A** ELISA(M2e) at 1/80 serum dilution (except PBS1X. 1/10 serum dilution) and **B** ELISA(Cell-M2) at 1/100 serum dilution. Results were expressed as OD values at a 1/100 serum dilution. **C** The ratios between ELISA(cM2) and ELISA(M2e) are indicated for each mouse. Data were log transformed and tested by **A and B** Mann-Whitney test or **C** paired t-test. Horizontal bars represent the geometric mean and vertical bars indicate the standard deviation of the geometric mean. P-values are given by *p<0.05. **p<0.01. ***p<0.001 and ****p<0.0001.

It is worth noting that M2 is *in vivo* primarily expressed on the surface of infected cells during influenza infection [5]. The recognition of infected cells by anti-M2e antibodies has been described as a key for flu protection. Therefore, ELISA(cM2) is more reliable in detecting anti-M2e mediating protection compared to ELISA(M2e). Indeed, free M2e peptides may display a large range of epitopes, which are not present in the M2 native protein. More critically, free M2e does not mimic the native M2e conformation as it appears during cellular M2 expression. ELISA(cM2) was performed with the same sera previously analyzed by ELISA(M2e) (Figure 3B). It appears that both free M2e and NLC-M2e induced comparable anti-cM2 antibody responses (average OD of 0.18 vs. 0.23, respectively, Figure 3B).

While antibody titer is an important parameter of the immune response, it is crucial to focus the antibody response against the target of interest for efficacy. Indeed, directing the immune response to a specific epitope ensures that the immune system concentrates its resources on optimizing affinity for that target, leading to more potent and specific antibodies [17]. Then, we examined the ratio of anti-cM2 antibodies to anti-M2e antibodies to assess how of both immunogens focus the antibody response to native epitopes expressed by cM2. Immunization with NLC-M2e resulted in a ∼6.7-fold higher ratio of cM2-specific antibodies compared to immunization with free M2e (0.15 vs. 1.0, respectively; p=0,03, Figure 3C). This observation highlights that the vectorization of M2e by NLC improves the specificity of antibody responses towards M2-expressing cells.

### A density of ∼40 M2e/NLC is sufficient for an optimal anti-M2e immune response

NLC-M2e formulations with different M2e densities (Table S2 and Figure 4) were used for immunizations with a constant NLC dose (0.5-0.6 × 10¹³ NLC). The results presented in Figure 4A indicated that higher M2e densities on the NLC surface led to stronger anti-M2e IgG responses (0.22 µg mL^-1^ for “Low”, 1.1 µg mL^-1^ for “Medium”, and µg mL^-1^ for “High”, with a statistically significant difference between “Low” and “High” with p=0.02, Figure 4A). An additional immunization was performed with a three times higher NLC-M2e dose (1.5 × 10¹³ NLC) than the initial dose using the “Medium” M2e density, thus varying both M2e and NLC doses. This was done in a separate immunization campaign with a different batch, where the slight difference in density between the two NLC-M2e “Medium” batches (47 M2e/NLC vs. 41 M2e/NLC) was considered negligible. This 3-fold increase in NLC-M2e dose resulted in a 3.8-fold increase in anti-cM2 IgG response compared to the lower dose of NLC-M2e “Medium” (4.3 µg mL^-1^ vs. 1.1 µg mL^-1^, Figure 4A), although this difference was not statistically significant. Interestingly, the higher dose of NLC-M2e “Medium” induced a similar response to that of the lower dose of NLC-M2e “High” (4.3 µg mL^-1^ vs. 3.2 µg mL^-1^, Figure 4A), despite the lower M2e dose (4.4 µg M2e vs. 2.8 µg M2e, Figure 4A). This trend suggests that above ∼40 M2e/NLC, the M2e dose has a greater impact on anti-cM2 IgG response than the M2e/NLC density

**Figure 4.**
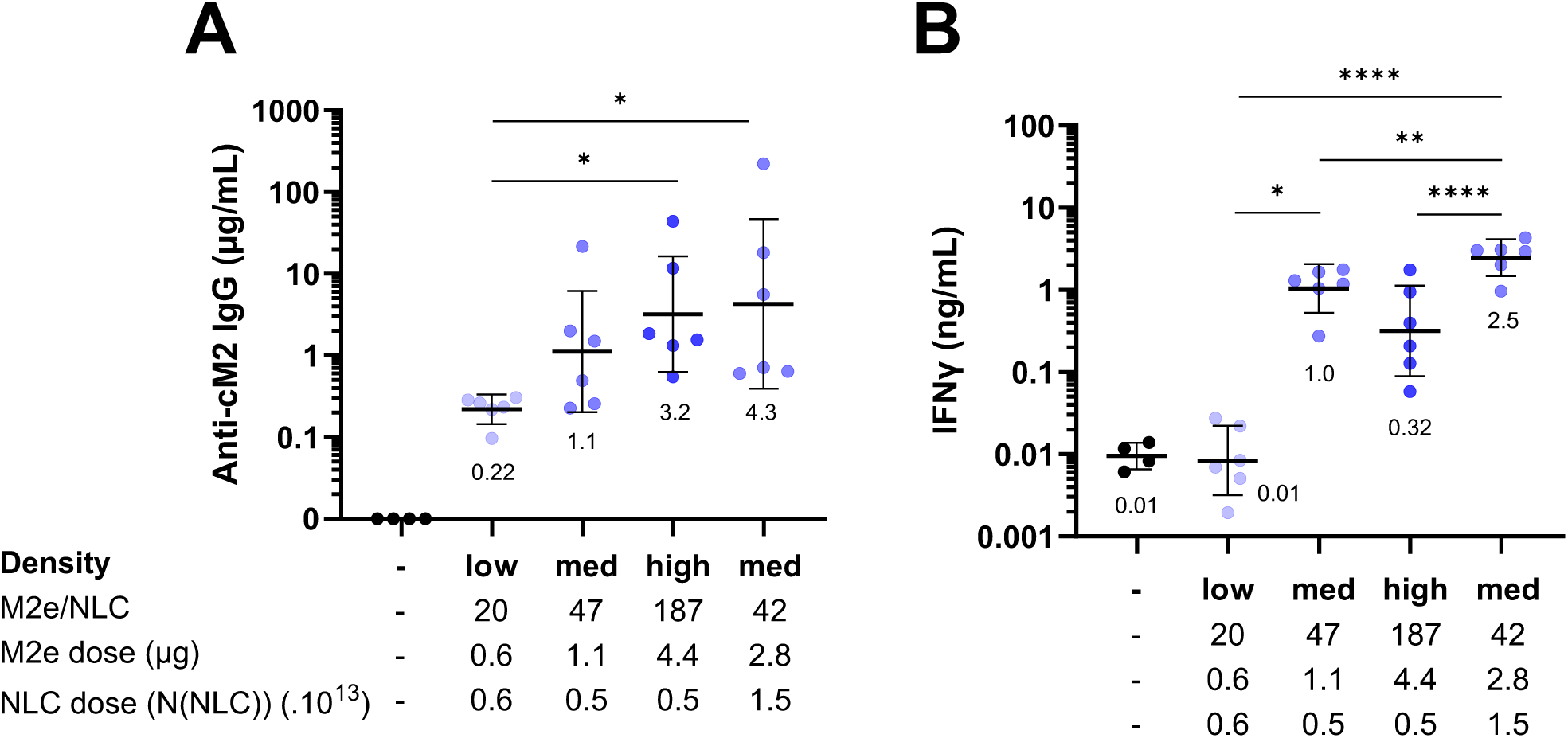
M2e dose rather than density increase allow a slight boost of anti-M2e response. BALB/c mice were immunized and double boosted with PBS 1x or NLC-M2e at two different NLC doses and/or with NLC-M2e of different densities. NLC-M2e were co-administered with a constant NLC(+)/CpG dose. Response was evaluated by **A** cM2 specific antibodies in the serum by ELISA(cM2) and by **B** IFNy production in the supernatant by ELISA of isolated splenocytes from immunized mice restimulated with M2e peptide. Data were **A** tested by Kruskal-Walis test followed by Dunn’s test or **B** tested by ordinary one-way ANOVA followed by Holm-Šídák’s multiple comparisons test. All the mice were not immunized and analyzed at the same time. Horizontal bars represent the geometric mean and vertical bars indicate the standard deviation of the geometric mean. P-values are given by *p<0.05. **p<0.01. ***p<0.001 and ****p<0.0001

Regarding cellular responses, monitored by IFNγ production of, NLC-M2e “Low” elicited no specific response to M2e, comparable to the negative control (0.01 ng mL^-1^ in both cases, Figure 4B). In contrast, other NLC-M2e immunizations induced significantly higher responses, with no statistically significant differences between NLC-M2e “Medium” and NLC-M2e “High” (1.0 ng mL^-1^ vs. 0.32 ng mL^-1^, p=0.38, Figure 4B). Increasing the NLC-M2e “Medium” dose led to a slight increase in IFNγ response (1.0 ng mL^-1^ vs. 2.5 ng mL^-1^, p=0.18, Figure 4B). Remarkably, the 1.5 × 10¹³ NLC-M2e “Medium” dose induced a statistically significant higher response than the 0.5 × 10¹³ NLC-M2e “High” dose (2.5 ng mL^-1^ vs. 0.32 ng mL^-1^, p=0.02, Figure 4B), despite having a lower M2e dose (2.8 µg M2e vs 4.4 µg M2e, Figure 4B).

Collectively, these data highlight that ∼40 M2e/NLC is sufficient for an optimal anti-M2e response, and that beyond this density, the M2e dose becomes more crucial.

### Enhancement of NLC-M2e Immune Response

To improve the NLC-M2e “Medium” formulation, we introduced a flexible linker composed of five amino acids (GGGGS) between the azide function and M2e (M2e-spacer-N_3_, Table S1). This modification was hypothesized to enhance the antibody response by increasing the spatial separation of M2e from the aromatic rings surrounding the cyclooctyne group (derived from the DBCO moiety) and by reducing steric hindrance imposed by the NLC. Immunization at a constant M2e dose elicited an 8.2-fold higher humoral response with NLC-spacer-M2e than with NLC-M2e (5.3 vs. 0.6 µg mL^-1^ anti-cM2 IgG, p=0.048, Figure 5A). The slight increase in NLC dose (1.4 × 10¹³ NLC-spacer-M2e compared to 1.1 × 10¹³ NLC-M2e) due to the difference in peptide density was insufficient to account for the significant increase in humoral response, which can be attributed to the spacer addition.

**Figure 5.**
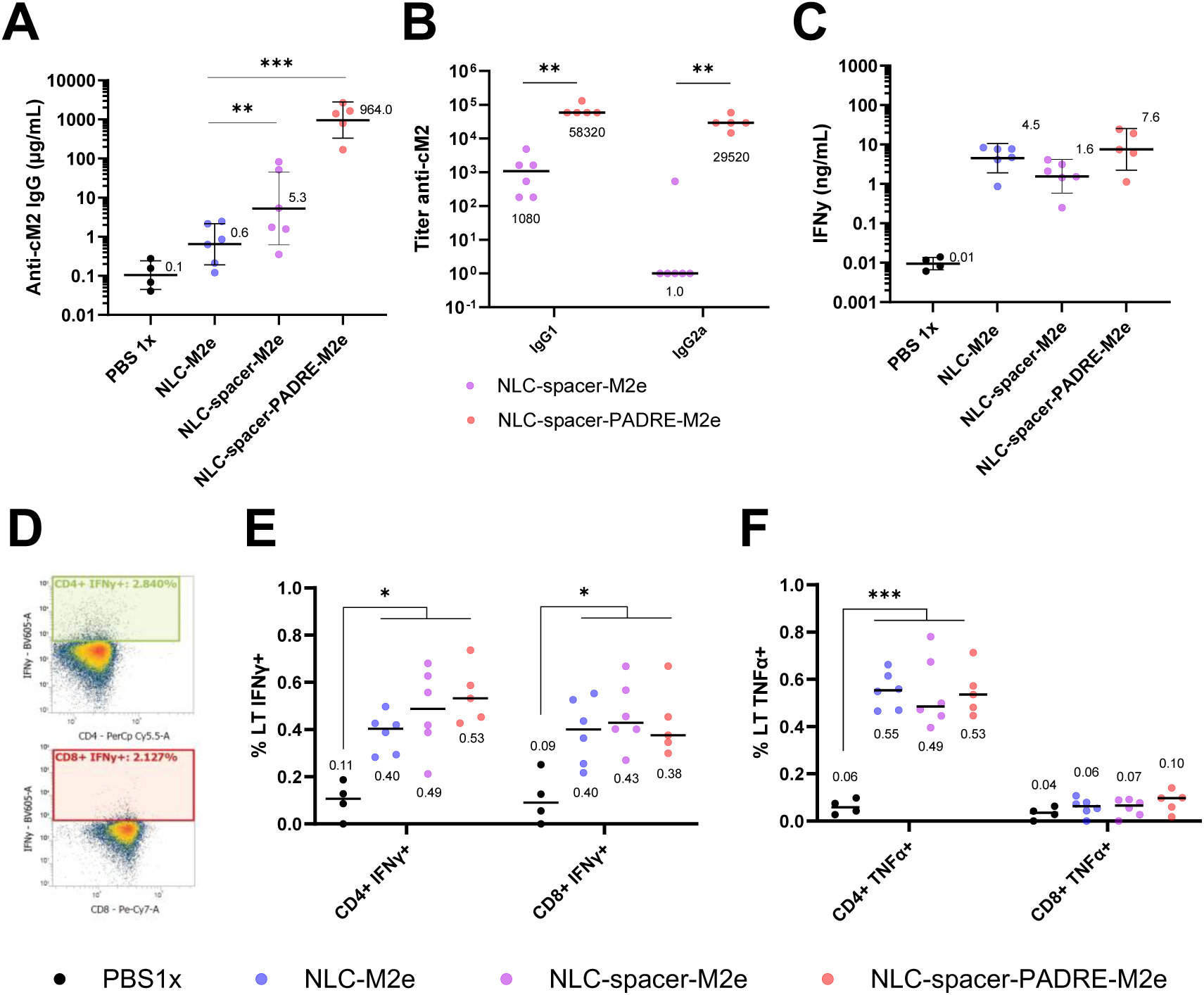
Insertion of a spacer and of PADRE peptide in NLC-M2e construction both allow massive augmentation of the anti-M2e antibody response. BALB/c mice were immunized and double boosted with either PBS 1x, NLC-M2e, NLC-spacer-M2e or NLC-spacer-PADRE-M2e. Corresponding formulation contained a dose of 1.9µg M2e and was co-administered with NLC(+)/CpG. Response was evaluated by **A** cM2 specific antibodies in the serum by cell-ELISA, by **B** IgG1 and IgG2a titer, by **C** IFNγ production in the supernatant by ELISA of isolated splenocytes from immunized mice restimulated with M2e peptide, by **D** the percentage of T lymphocytes intracellularly expressing either **E** IFNγ or **F** TNFα after restimulation with M2e peptide. Data were **A** log transformed and tested by ordinary one-way ANOVA followed by Holm-Šídák’s multiple comparisons test, **B** not transformed and tested by Mann-Withney tests, **C** log transformed and tested by Kruskal-Walis test followed by Dunn’s test, **E and F** not transformed and tested by ordinary one-way ANOVA followed by Holm-Šídák’s multiple comparisons test. Horizontal bars represent the geometric mean and vertical bars indicate the standard deviation of the geometric mean. P-values are given by *p<0.05. **p<0.01. ***p<0.001 and ****p<0.0001

Next, we fused the M2e-spacer-N_3_ to a universal class II MHC peptide named PADRE as a help for T cell responses [14] creating the M2e-PADRE-spacer-N_3_ peptide (Table S1). Strikingly, immunization with NLC-spacer-PADRE-M2e induced a 182-fold higher anti-cM2 IgG response compared to NLC-spacer-M2e (964 vs. 5.3 µg mL^-1^ anti-cM2 IgG, p=<0.0001, Figure 5A).

We then analyzed the quality of the antibody response, in terms of Ig subclasses, by measuring IgG1 and IgG2a titers. NLC-spacer-M2e elicited an almost exclusive IgG1 response (titer of 1080, Figure 5B, while the addition of PADRE produced a more balanced response between IgG1 and IgG2a (titers of 58,320 and 29,520, respectively, Figure 5B). It is important to note that the affinity of anti-M2e antibodies for cM2 was only moderately modified by the addition of PADRE (factor of 3, Figure S4). Interestingly, both polyclonal responses generated by NLC-spacer-M2e and NLC-spacer-PADRE-M2e demonstrated superior affinity compared to the anti-M2e monoclonal antibody 14C2, which served as a reference.

We also evaluated the cellular immune response to the different formulations. The addition of the spacer or PADRE did not significantly affect IFNγ production compared to NLC-M2e (4.5 ng mL^-1^, 1.6 ng mL^-1^, and 7.6 ng mL^-1^ of IFNγ, respectively, Figure 5C). Flow cytometry analysis revealed that for all formulations, IFNγ was produced by both CD4+ and CD8+ T cells (Figure 5D and Figure 5E). Additionally, TNFα production was found exclusively in CD4+ T cells (Figure 5F). Neither the addition of the spacer nor the PADRE peptide significantly increased TNFα production.

## DISCUSSION

Nanostructured Lipid Carriers (NLCs) are lipid nanoparticles (LNPs) characterized by a solid lipidic core, which strongly enhances their stability in solution. By modifying their surface chemistry or charge, antigens and nucleotides can be effectively vectorized on their surface. Initial studies have demonstrated their efficacy as vaccine vectors in delivering whole protein antigens [9, 10] or mRNA encoding antigens [18].

In this study, our objective was to expand NLC application to small antigens such as peptides, which are extensively used in vaccines [19] but still lack licensed prophylactic vaccines. We selected the M2e peptide from the influenza virus as our antigenic peptide due to its potential as a universal flu vaccine candidate [5]. However, its low immunogenicity currently hinders its further use in vaccination. Therefore, eliciting robust immune responses against the M2e peptide remains a significant challenge.

The native orientation of M2e on infected cells exposes its N-terminal portion, which has been shown to generate a stronger antibody response than the C-terminal portion [20, 21]. Additionally, M2e is a short peptide with few epitopes, making it crucial to preserve key recognition sites, particularly cysteines C17 and C19. These cysteines allow the formation of a conformational epitope critical for protection [22]. Therefore, the widely used thioether/maleimide reaction, which can react with internal cysteines, is unsuitable for optimal control of M2e orientation after conjugation. Binding M2e via its C-terminal portion without affecting internal amino acids would maintain the cysteines free, thus preserving potential internal epitopes. To address this, we selected Click Chemistry, specifically the SPAAC reaction, which offers the key advantage of being bioorthogonal, ensuring preservation of M2e structure [16]. It requires two chemical groups: an azide (N_3_) and dibenzoazacyclooctynes (DBCO). N_3_ insertion into the C-terminal end of M2e was achieved during peptide synthesis using a modified amino acid (Orn-N_3_), making it a cost-effective and reliable approach. On the other hand, DBCO was introduced at the surface of NLC (NLC-DBCO) using SA-PEG_100_-DBCO during NLC formulation. NLC-DBCO enabled conjugation of M2e-N_3_ with a maximum conjugation rate of 63.6%, comparable to values reported in the literature [23]. NLC-DBCO showed only slight reactivity loss over time (Table S4) and remained usable for up to three months in 1x PBS. For instance, NLC-DBCO was more stable than NLC-Maleimide formulated in similar conditions [24], which loses 30% of reactivity within one month, whereas NLC-DBCO showed only an 18.7% loss in almost three months. These results highlight the robustness of the SPAAC reaction for M2e conjugation.

We then tested our NLC-M2e construct *in vivo*. Immunization with NLC-M2e compared to free M2e allowed the anti-M2e humoral response to focus on cM2 recognition (Figure 3, Figure S3). Conserving M2e key epitopes ensures preservation of antigen-specific immune responses, including humoral-specific responses. This is particularly important, as studies have demonstrated that anti-M2e antibodies mediate protection by opsonizing influenza-infected cells expressing M2e [4]. In line with this mechanism, other studies have shown that detection of the antibody response by cells expressing M2 (cM2) correlates with protection [25, 26]. After being opsonized by M2e-specific antibodies, infected cells are destroyed through antibody-dependent cellular phagocytosis (ADCP) by macrophages or antibody-dependent cell-mediated cytotoxicity (ADCC) by natural killer cells [27–29], two mechanisms requiring strong support from Th1 cells, particularly through IFNγ signaling. Increasing the magnitude of both humoral and cellular specific anti-M2e responses was therefore mandatory.

To enhance the strength of the immune response, it has been reported that antigen and/or NLC dosage [30] as well as antigen density on the NLC surface might be critical [8]. Our study showed that a minimum M2e density per NLC (∼40 M2e/NLC), was necessary for eliciting robust humoral and cellular responses (Figure 4). Increasing M2e density beyond this point did not significantly boost the humoral or cellular response. Above 40 M2e/NLC, the key parameter to increase is the global M2e dose. For instance, previous studies showed that higher density of M2e peptide per nanoparticle enhanced the immune response [31, 32]. However, a more in-depth study using PapMV NPs demonstrated a plateau in the humoral response at higher antigenic densities [33], which is consistent with our findings.

Another way to enhance immune response toward M2e was to ensure that its accessibility at the NLC-M2e surface was optimal. We hypothesized that the hydrophobic moiety remaining from the DBCO moiety at the C-terminal end of M2e sequence may hinder antibody binding to key M2e epitopes. Inserting a peptide spacer between the C-terminal end of M2e and the azide group improved antibody response by 8.8-fold without affecting cellular response (Figure 5A). This may be due to the spacer alleviating steric hindrance, thus allowing freedom for epitope formation. Indeed, structural studies of M2e in complex with the M2e-specific monoclonal antibody mAb65 revealed that M2e adopted a loop-like conformation during interaction, reducing the distance between its C-terminal and the rest of the peptide [34], supporting our hypothesis.

The last mechanism explored to improve the anti-M2e response was the CD4+ T lymphocyte support for anti-M2e antibodies production. It has been established that the number of T-cell epitopes in M2e might be suboptimal for achieving high cellular responses across a wide range of MHC molecules, as only a single T-human epitope was clearly identified previously [35]. Therefore, enhancing T-helper (Th) cell responses to boost the anti-M2e antibody responses is suitable. To test this possibility, we fused M2e with T helper antigen: the PADRE peptide. PADRE has the ability to be presented by both human and mouse MHC-II molecules, HLA-DR and H-2 IA, making it useful in preclinical demonstration and clinical applications as a universal helper epitope [36]. PADRE-specific Th cells only enhance antibody responses to linked epitopes, reducing the risk of off-target responses. PADRE has demonstrated a strong safety profile in phase I clinical trials [37] and was found to be more effective than most other universal epitopes, stimulating CD4+ T cells 100 times more potently than tetanus toxin epitopes [14]. Our results showed that incorporating PADRE as a helper epitope supported M2e-specific B cell responses, leading to a dramatic ∼180-fold increase in anti-cM2 antibody response compared to NLC-spacer-M2e (Figure 5A). This finding is in line with the previously mentioned studies and supports the rational of introducing PADRE in our construct. Notably, the efficacy of PADRE peptide in humans increases the likelihood of successful translation to clinical settings. When combined with the efficient antigen presentation enabled by NLC vectorization, the NLC-spacer-PADRE-M2e formulation holds strong potential to overcome the limitations observed with earlier M2e-based vaccine candidates [5].

We also assessed the quality of T-lymphocyte response. Our data revealed both CD4+ and CD8+ IFNγ T cell responses following M2e immunization, although only CD4+ T cell epitopes for M2e have been described in BALB/c mice [35, 38]. The role of CD8+ T cells in M2e-based immunity remains unclear, but evidence points to their potential involvement in protection [39]. The presence of both CD4+ and CD8+ T-lymphocyte populations might help for *in vivo* protection against flu. In addition, the Th1 phenotype induced by our formulation suggests robust opsonization of infected cells via IgG2a, facilitating ADCP and ADCC. Importantly, all mice immunized with NLC-spacer-PADRE-M2e surpassed the threshold IgG2a titer required for protection identified by Fiers et al. [40] and Rappazzo et al. [41] (Figure S5).

Lastly, the NLC-spacer-PADRE-M2e formulation might be mixed with HA or NA entire antigen or added to current flu vaccines formulations, potentially increasing protection across various influenza subtypes [42–46]. Future studies should evaluate the efficacy of NLC-spacer-PADRE-M2e *in vivo*. Protecting mice from lethal influenza infection using this vaccine would confirm its potential as a universal flu vaccine candidate, capable of providing broad, long-lasting immunity against both seasonal and pandemic strains of influenza.

## CONCLUSION

In conclusion, our study demonstrates the potential of NLC as a versatile vaccine delivery system for poorly immunogenic antigens like the M2e peptide, a key target for universal influenza vaccines. Through efficient biorthogonal SPAAC chemistry, we achieved stable and high-density conjugation of M2e to NLC, preserving crucial epitopes and enabling robust antigen presentation. The addition of a flexible spacer significantly enhanced the humoral immune response, likely by improving B cell recognition. Moreover, the incorporation of the PADRE peptide further amplified the anti-M2e antibody response by stimulating helper T cell activity, achieving a 180-fold increase in IgG titers. While M2e-specific CD4+ IFNγ T cells were anticipated, we also observed the unexpected involvement of CD8+ T cells, suggesting a broader cellular response than previously described in the literature. The robust Th1-type response, characterized by high IFNγ levels and IgG2a titers, further supports the potential protective efficacy of this vaccine platform, particularly through mechanisms such as ADCP.

This work underscores the flexibility of the NLC platform for multi-antigen delivery, and the addition of universal T helper epitopes like PADRE opens the door for further refinement of immune responses. The ability to modulate the density, conjugation, and helper peptide inclusion offers a pathway to design more effective and targeted vaccines.

Future research should focus on *in vivo* challenge studies to confirm the protective efficacy of this candidate and explore broader immune signatures. Together, these findings provide a strong foundation for the continued development of NLC-based vaccines targeting M2e and other antigens of interest.

## AUTHORS’ CONTRIBUTIONS

Louis Bourlon: Investigation; Formal analysis, Writing – original draft;

Carole Fournier: Methodology; Formal analysis, Writing – review and editing;

Antoine Hoang: Methodology; Formal analysis;

Christine Charrat: Methodology; Formal analysis;

Alban Lepetitcolin: Supervision, Project administration

Jessica Morlieras: Investigation, Data curation, Formal analysis Funding acquisition; Writing – review and editing;

Fabrice Navarro: Conceptualization, Formal analysis, Funding acquisition, Writing – review and editing;

Patrice N Marche: Conceptualization, Formal analysis, Funding acquisition, Writing – review and editing;

These authors contributed equally: Patrice N. Marche, Fabrice P. Navarro

## ACKNOWLEDGEMENTS

We thank the staff of the core facilities of the IAB especially Mylène Pezet, for her assistance with flow cytometry equipment, and the personal of the animal facility for animal care. We thank Marie Escudé (CEA) for her help for NLC formulation, Jérôme Jude (Sanofi) for the help for AAA-UPLC, Carole Borovsek (Sanofi) for the help for ELISA(NLC-M2e) set-up. We also thank Cédric Charretier (Sanofi) and Matthieu Varache (Sanofi) for their advices. The authors also thank Jean-Sébastien Bolduc for critical review and proofreading of the manuscript.

## GRANTS AND SUPPORTS

The project was supported in part by the French Ministry of Higher Education, research and innovation [LipiVAC, COROL project, funding reference N◦ 2102992411]. The project used Microcell facility of IAB which is part of the IBISA-ISdV platform, member of the national infrastructure FranceBioImaging supported by the French National Research Agency (ANR-10-INBS-04).

L.B was supported by a specific doctoral grant from the French Association Nationale Recherche Technologie (CIFRE 2020/0228).

C.F was supported by a fellowship from INSERM Transfert (Proof of Concept 2021) and by the foundation “FINOVI” (2022).

## GLOSSARY

Abbreviation: Signification
NLC: Nanostructured Lipid Carrier
M2e: Matrix Protein 2 Ectodomain
HA: Hemagglutinin
LNP: Lipid Nanoparticle
PEG: Polyethylene Glycol
SPAAC: Strain-Promoted Azide-Alkyne Cycloaddition
IFNγ: Interferon gamma
DOPE: 1,2-Dioleoyl-sn-glycero-3-phosphoethanolamine
DMSO: Dimethyl sulfoxide
DLS: Dynamic Light Scattering
NTA: Nanoparticle Tracking Analysis
TLC: Thin Layer Chromatography
SEC: Size Exclusion Chromatography
AAA: Amino Acid Analysis
UPLC: Ultra Performance Liquid Chromatography
_CpG_: Cytosine-phosphate-Guanine (DNA motif used as adjuvant)
ELISA: Enzyme-Linked Immunosorbent Assay
OD: Optical Density
TMB: Tetramethylbenzidine
UTR: Untranslated Region
PADRE: Pan-DR Epitope
TNFα: Tumor Necrosis Factor alpha
FMO: Fluorescence Minus One
ADCC: Antibody-Dependent Cell-mediated Cytotoxicity
ADCP: Antibody-Dependent Cellular Phagocytosis
Th1: T-helper 1 cell type
FCSB: Flow Cytometry Staining Buffer

