## Supplementals for "Nanostructured lipid carriers overcome the low immunogenicity of M2e peptide via surface click chemistry conjugation, improving the anti-M2e antibody response"

### Slide 1
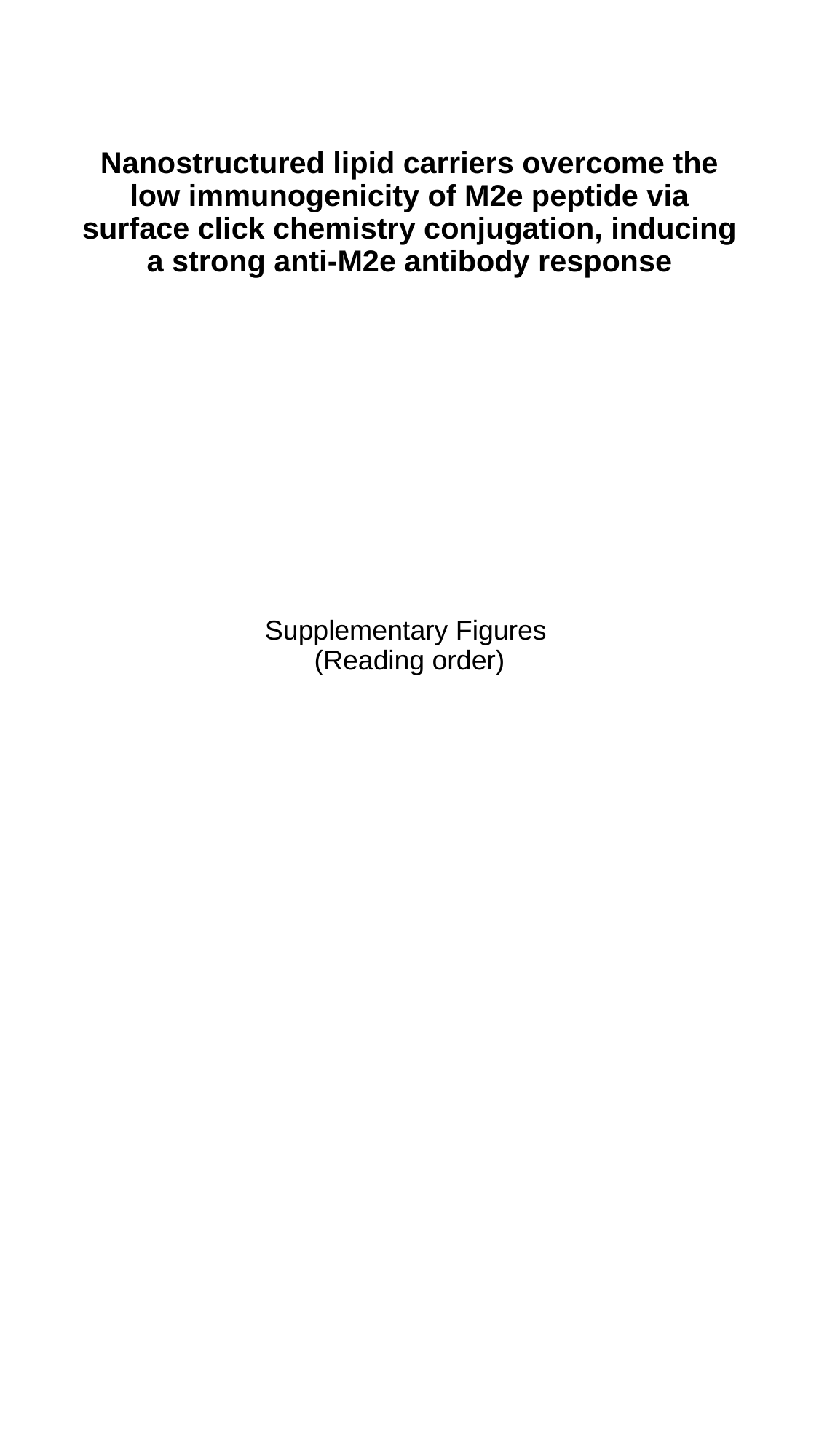

#### Nanostructured lipid carriers overcome the low immunogenicity of M2e peptide via surface click chemistry conjugation, inducing a strong anti-M2e antibody response
Supplementary Figures
(Reading order)

### Slide 2
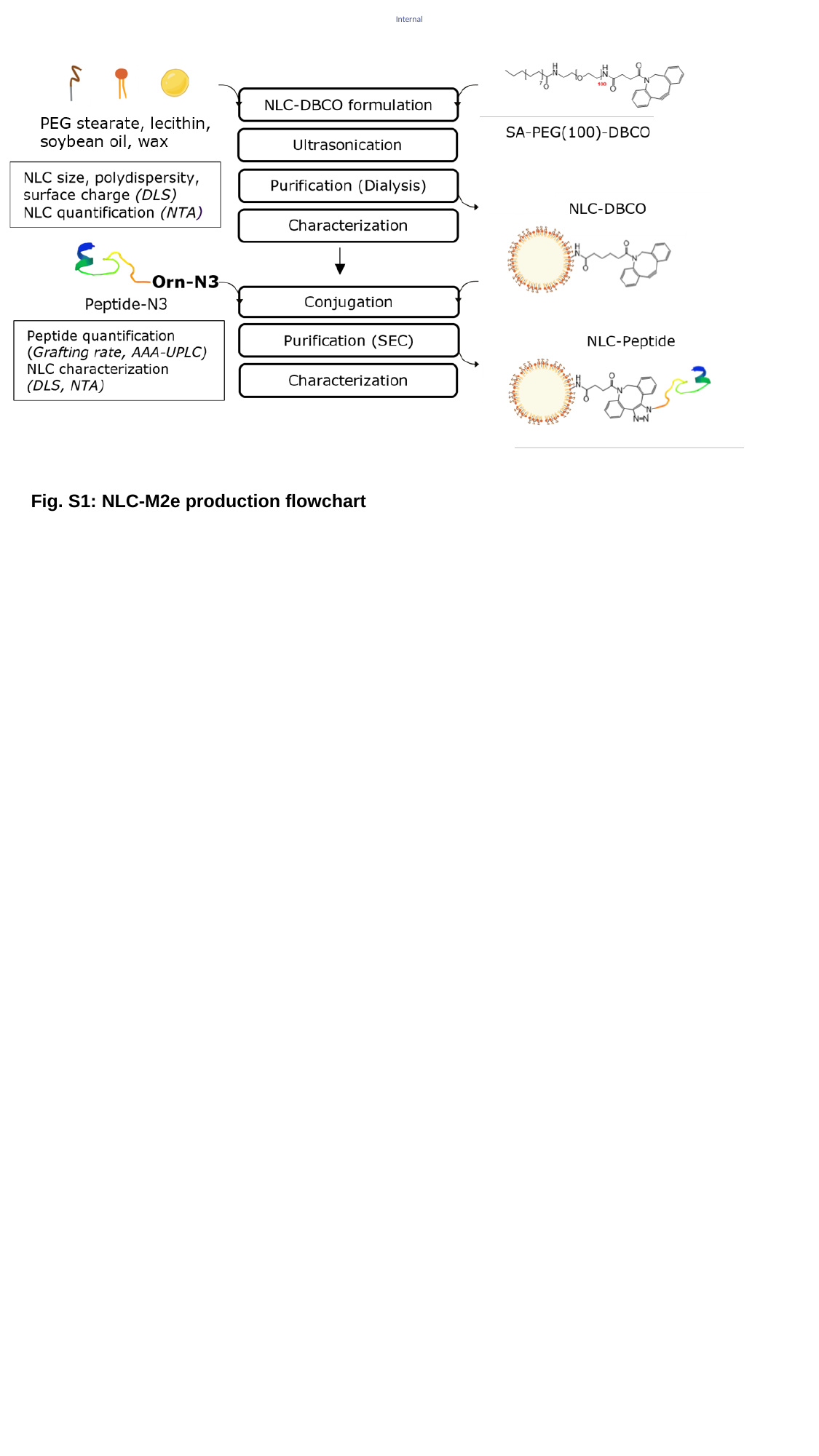

Fig. S1: NLC-M2e production flowchart

### Slide 3
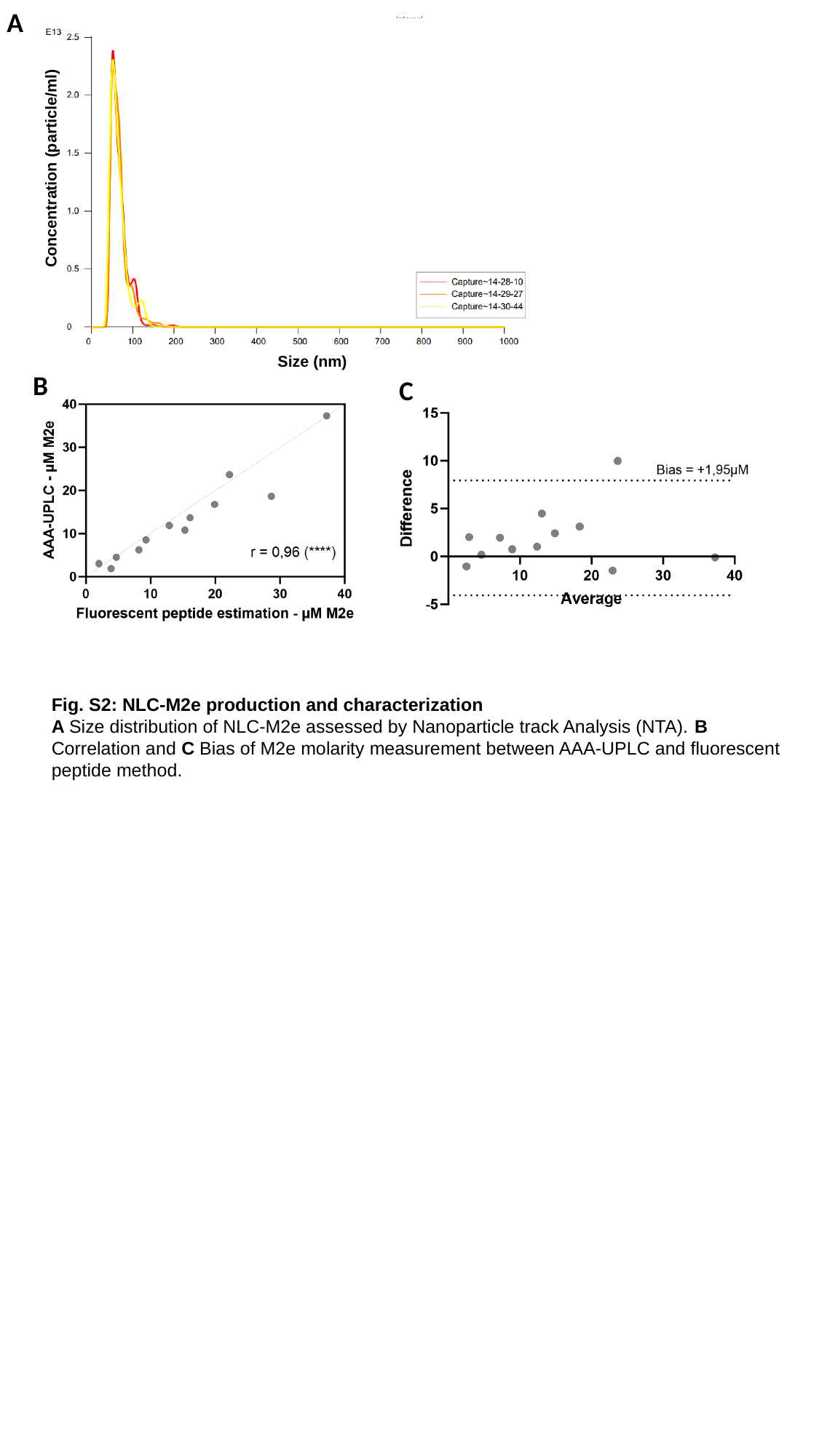

A
B
C
Fig. S2: NLC-M2e production and characterization
A Size distribution of NLC-M2e assessed by Nanoparticle track Analysis (NTA). B Correlation and C Bias of M2e molarity measurement between AAA-UPLC and fluorescent peptide method.

### Slide 4
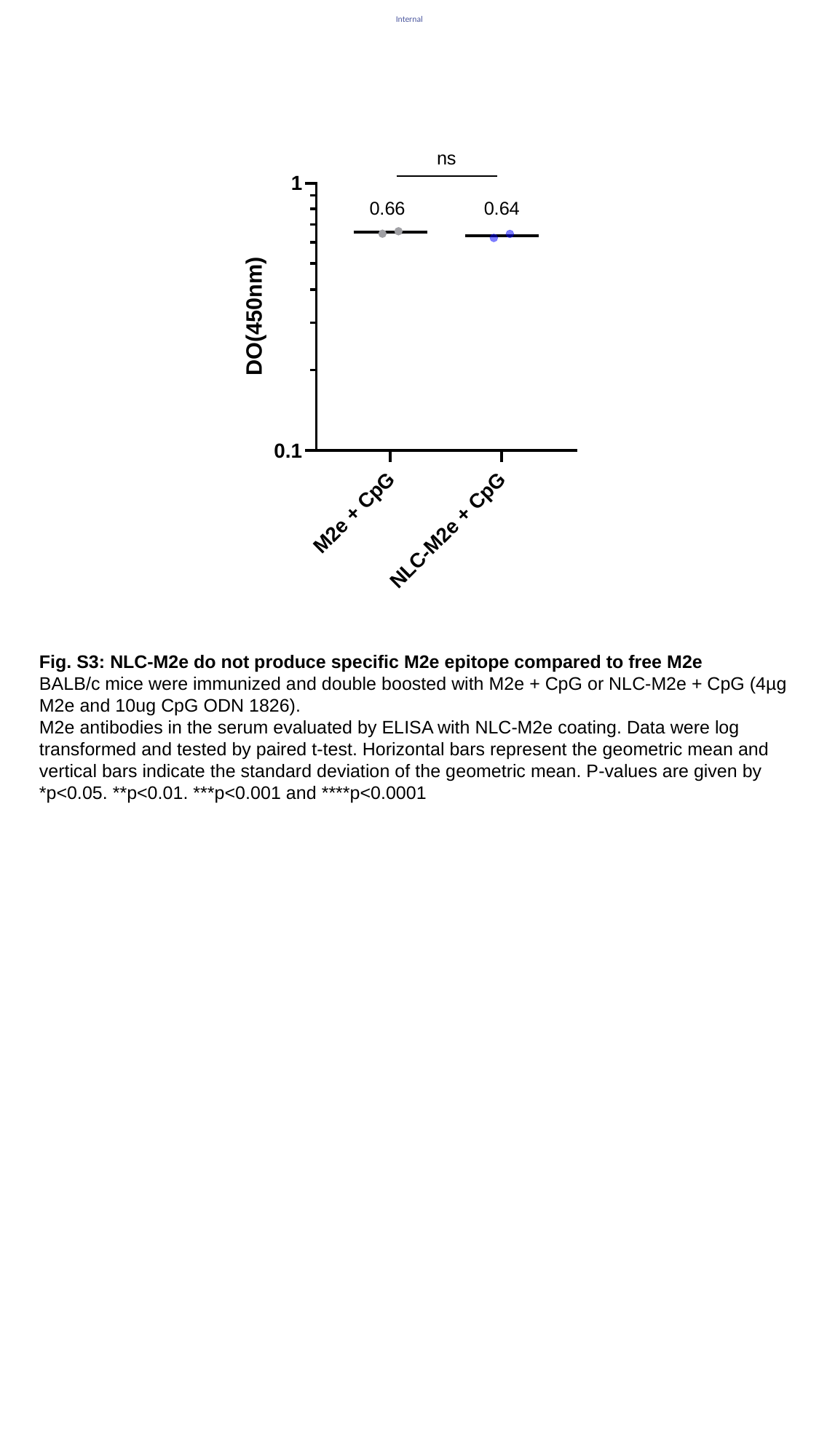

Fig. S3: NLC-M2e do not produce specific M2e epitope compared to free M2e
BALB/c mice were immunized and double boosted with M2e + CpG or NLC-M2e + CpG (4µg M2e and 10ug CpG ODN 1826).
M2e antibodies in the serum evaluated by ELISA with NLC-M2e coating. Data were log transformed and tested by paired t-test. Horizontal bars represent the geometric mean and vertical bars indicate the standard deviation of the geometric mean. P-values are given by *p<0.05. **p<0.01. ***p<0.001 and ****p<0.0001

### Slide 5
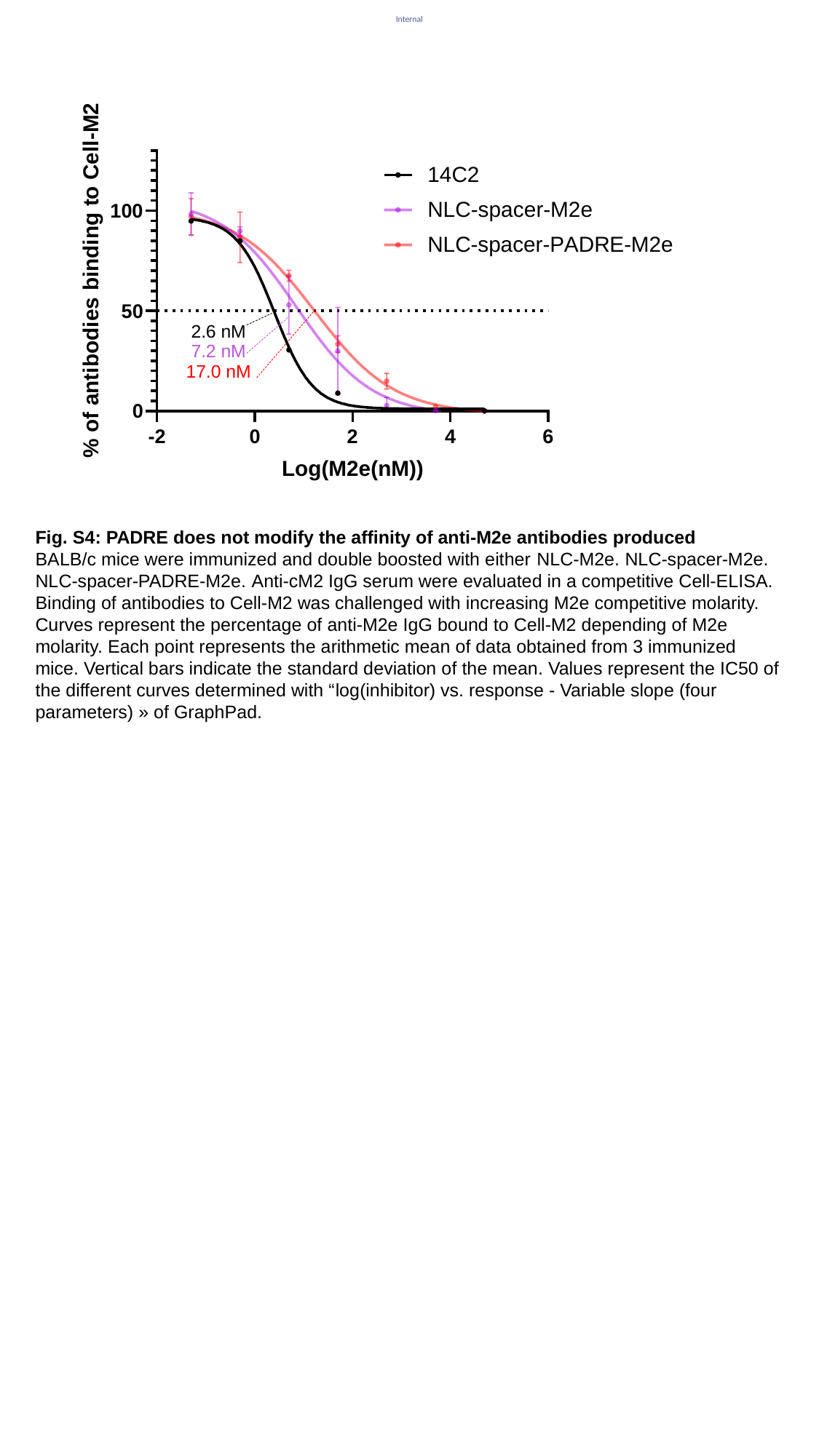

Fig. S4: PADRE does not modify the affinity of anti-M2e antibodies produced
BALB/c mice were immunized and double boosted with either NLC-M2e. NLC-spacer-M2e. NLC-spacer-PADRE-M2e. Anti-cM2 IgG serum were evaluated in a competitive Cell-ELISA. Binding of antibodies to Cell-M2 was challenged with increasing M2e competitive molarity. Curves represent the percentage of anti-M2e IgG bound to Cell-M2 depending of M2e molarity. Each point represents the arithmetic mean of data obtained from 3 immunized mice. Vertical bars indicate the standard deviation of the mean. Values represent the IC50 of the different curves determined with “log(inhibitor) vs. response - Variable slope (four parameters) » of GraphPad.

### Slide 6
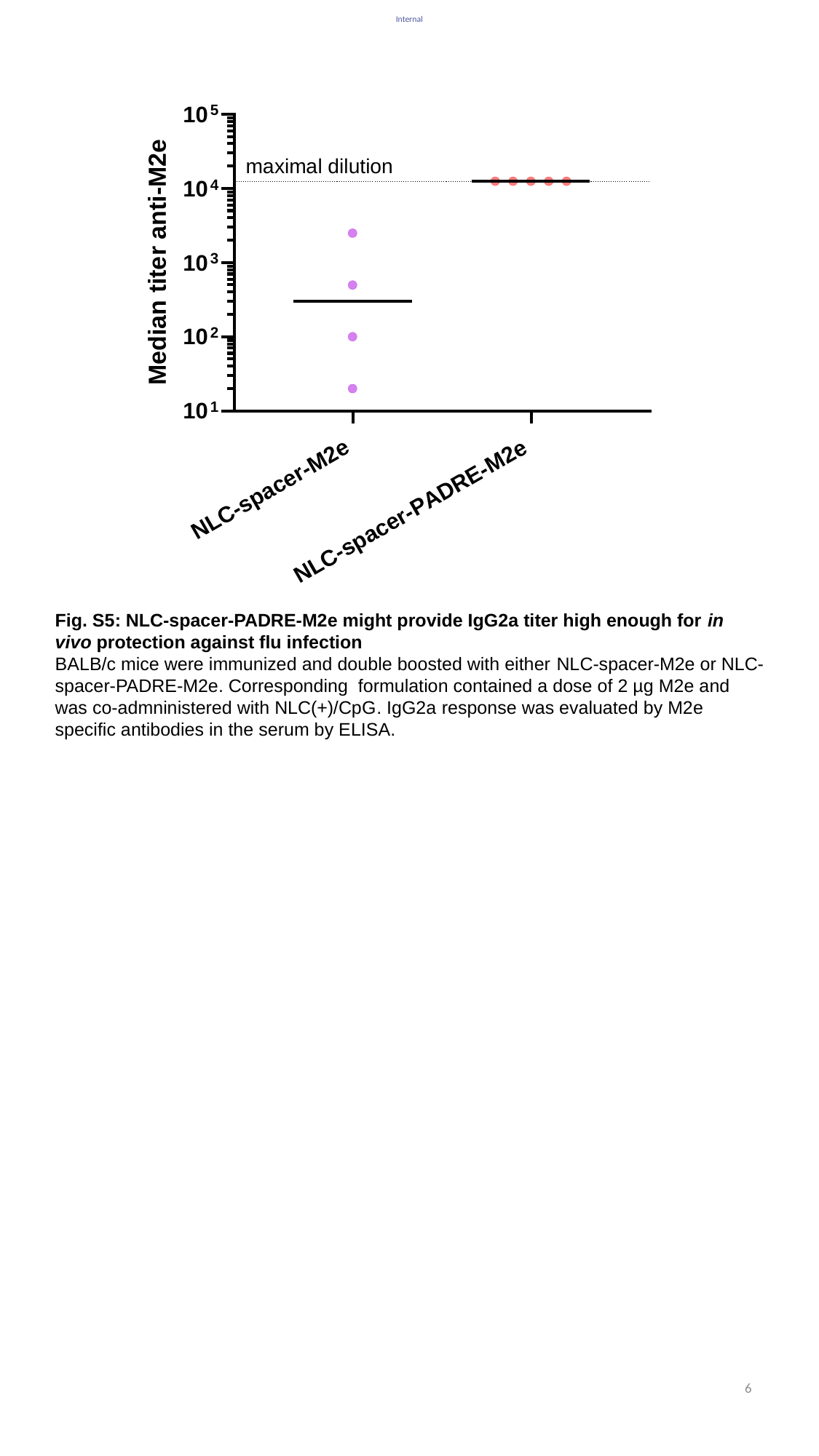

Fig. S5: NLC-spacer-PADRE-M2e might provide IgG2a titer high enough for in vivo protection against flu infection
BALB/c mice were immunized and double boosted with either NLC-spacer-M2e or NLC-spacer-PADRE-M2e. Corresponding formulation contained a dose of 2 µg M2e and was co-admninistered with NLC(+)/CpG. IgG2a response was evaluated by M2e specific antibodies in the serum by ELISA.
6
